# Zebrafish cooperate while inspecting predators: experimental evidence for conditional approach

**DOI:** 10.1101/814434

**Authors:** Ana Flávia Nogueira Pimentel, Monica Gomes Lima-Maximino, Marta Candeias Soares, Caio Maximino

## Abstract

Different fish species employ a conditional approach strategy during predator inspection; the risk of approaching a predator is distributed across all inspectors, but is not shared with the animals which keep its distance. Zebrafish, a highly social fish, is increasingly being used in behavioural neuroscience, but it is not known whether it displays conditional approach. In the predator inspection task, animals are observed in a tank with a refuge in one extremity, and an animated image of a predator in the other extremity, with a mirror positioned in parallel to the tank, simulating a perfectly reciprocating conspecific. In Experiment 1, animals spent more time in an inspection zone when the image was turned on, but also displayed more erratic swimming, suggesting cooperation under fear. In Experiment 2, animals spent more time inspecting predators when the mirror was parallel to the tank (“cooperating mirror”) than when the mirror was in an angle (“defecting mirror”), suggesting retaliatory behaviour; in both conditions, animals displayed more freezing and erratic swimming. In Experiment 3, no changes in behaviour were observed, regardless of mirror position, when the image was turned off, suggesting that the choice of specific zones in Experiment 2 was not due to shoaling tendencies. These results suggest that predator inspection is associated with conditional approach, while at the same time inducing fear-like behaviour in the animal.

## 1. Introduction

While being widespread among animals, altruism is an adaptive/evolutionary conundrum, as it may involve reducing one’s own fitness in order to benefit another individual (Kropotkin, 1902). Different explanations have been given to this phenomenon, including kin selection (W. D. Hamilton, 1963, 1964) and reciprocal altruism (Trivers, 1971). Reciprocal altruism predicts that this type of cooperative behaviour only occurs in populations when it increases the likelihood of reciprocation at the receiving end – that is, the individual decreases its own fitness in the expectation that the other organism will act in a similar manner in a later time (Trivers, 1971). Game theoretical approaches suggest a “tit-for-tat” strategy, in which an individual cooperates in the first encounter and continues to cooperate as long as the other agent does as well (Trivers, 1971).

Experimental evidence of “tit-for-tat” has been hard to produce because the payoff (i.e., possible positive and negative outcomes) of cooperating vs. defecting is known, but the benefits (and risks) cooperating are usually not known. Nonetheless, “reciprocal altruism-like” behaviour has been observed in many species (Bshary & Oliveira, 2015). An interesting example is conditional approach, a strategy used by social fish when approaching a potential predator, leaving the shoal to obtain information on putative threat and/or try to intimidate the dangerous predator (Dugatkin, 2013). For two interacting fish approaching a predator, defecting (staying behind) is beneficial because the defecting individual decreases its risk but increases its payoff (e.g., information gathered) by watching the fate of the other fish. Conditional approach predicts that inspecting fish should immediately retaliate defecting conspecifics by also defecting (Bshary & Oliveira, 2015; Dugatkin, 1988; Pimentel et al., 2019). The strategy appears to function as an incentive for predator inspection, given that the risk of approaching it is distributed between all inspectors but not shared with those that remain at a distance. Since this behaviour implies costs to the individual(s) interacting with the predator, but contributes to the shoal’s fitness, it has been interpreted as cooperative-like and altruistic (Dugatkin, 1988, 1997; Dugatkin & Alfieri, 1991b, 1992; Pimentel et al., 2019). When choosing whether or not to inspect a predator, individuals leaving the shoal are exposed to increased predation risk (Dugatkin, 1992; Milinski et al., 1997), and therefore are expected to display threat-sensitive behaviours such as freezing and erratic swimming. In guppies, predator inspection is accompanied by increased freezing duration, suggesting that this is indeed the case (Pimentel et al., 2019).

In conditional approach, prey approaching predators inspect the predator at the first “move”, immediately head back away from the predator if the conspecific falls behind or disappears, and approach the predator again if the conspecific’s move of “swimming away and staying behind” is followed by “swim parallel” (Dugatkin, 1988). However, differently from tit-for-ta t(Axelrod & Hamilton, 1981), conditional approach does not require that the actual payoff matrix is known (Dugatkin, 1988). Conditional approach has been shown in sticklebacks (*Gasterosteus aculeatus* Linnaeus 1758) (Huntingford et al., 1994; Milinski, 1987) and guppies (*Poecilia reticulata* Peter 1859) (Dugatkin, 1988, 1991; Dugatkin & Alfieri, 1991b, 1992; Pimentel et al., 2019), but not for zebrafish, an important model organism in the neurosciences.

Zebrafish (*Danio rerio* Hamilton 1822) is a model organism that is widely used in developmental biology (Parichy, 2015) and which has been introduced in the field of behavioural neuroscience and behavioural ecology (Bonan & Norton, 2015; Stewart et al., 2015). This fish is also a good model organism for biological psychiatry (Stewart et al., 2015), due to the availability of behavioural bioassays and the advantages associated with being a model organism, including low cost of acquisition and maintenance, easy handling and housing, short lifespan, and easy reproduction in the laboratory. As a highly social animal that lives in groups with well-structured social relations – including marginalization, dominance hierarchies, and territoriality (Spence et al., 2008) –, zebrafish are also known to display predator inspection (Dugatkin et al., 2010; Pannia et al., 2014); however, it is not known whether zebrafish employ conditional approach when inspecting predators.

In addition to helping to better understand complex social behaviour in this species, here we aim to introduce new behavioural techniques that can also advance research in the field of neuropsychopharmacology, including the development of drugs to treat disorders of social behaviour (Soares et al., 2018). While the focus of this field on zebrafish social behaviour has relied mainly on shoaling, more complex behaviours, such as cooperative-like behaviour, could represent an important addition to the field. Our main aim was to investigate preliminary evidence of conditional approach in zebrafish in a predator inspection model, based in Milinski (1987) and Dugatkin (1988). In Experiment 1, a mirror was placed in parallel to a tank, and the animated image of a predator (Gangetic leaffish *Nandus nandus* Hamilton 1822): when the animation was turned on, animals spent more time in the inspection area than when it was turned off, suggesting that the presence of both a predator and a cooperating conspecific (mirror image) are necessary for inspection behaviour. In Experiment 2, the mirror was either placed in parallel (“cooperating mirror”) or in an angle of 45° with the side of the tank which was opposite to the predator (“defecting mirror”): animals then spent less time in the inspection zone during the defecting mirror condition. In Experiment 3, animals were exposed to conditions of Experiment 2, but the video was not turned on. Overall, animals displayed freezing and erratic swimming when the animation was visible, suggesting fear. This manuscript is the first complete report of all the studies performed to test the hypothesis that zebrafish display conditional approach during predator inspection. We report all data exclusions (if any), all manipulations, and all measures in the study.

## 2. Methods

### 2.1. Animals and housing

31 animals were used in Experiments 1 and 2 described below; data from one animal was excluded from each experiment due to poor health after the end of the experiments, which could impact the results. In Experiment 3, 32 animals were used. Thus, a total of 94 animals were used across all three experiments. Animals were bought from a commercial vendor, and arrived in the laboratory with an approximate age of 3 months (standard length = 13.2 ± 1.4 mm), and were quarantined for two weeks; the experiment began when animals had an approximate age of 4 months (standard length = 23.0 ± 3.2 mm). Animals were kept in mixed-sex tanks during acclimation, with an approximate sex ratio of 1:1 (confirmed by body morphology). Adult zebrafish from the wildtype strain (longfin phenotype) were used in the experiments. Outbred populations were used due to their increased genetic variability, decreasing the effects of random genetic drift which could lead to the development of uniquely heritable traits (Parra et al., 2009; Speedie & Gerlai, 2008). Thus, the animals used in the experiments are expected to better represent the natural populations in the wild. The breeder was licensed for aquaculture under Ibama’s (Instituto Brasileiro do Meio Ambiente e dos Recursos Naturais Renováveis) Resolution 95/1993. Animals were group-housed in 40 L tanks, with a maximum density of 25 fish per tank, for at least 2 weeks before experiments begun. Tanks were filled with non-chlorinated water at room temperature (28 °C) and a pH of 7.0-8.0 (coefficient of variation = 5.01%). Lighting was provided by fluorescent lamps in a cycle of 14-10 hours (LD), according to standards of care for zebrafish (Lawrence, 2007). Water quality parameters were as follows: pH 7.0-8.0; hardness 100-150 mg/L CaCO3; dissolved oxygen 7.5-8.0 mg/L; ammonia and nitrite < 0.001 ppm. All manipulations minimized their potential suffering of animals, and followed Brazilian legislation (Diretriz Brasileira Para o Cuidado e a Utilização de Animais Para Fins Científicos e Didáticos - DBCA. Anexo I. Peixes Mantidos Em Instalações de Instituições de Ensino Ou Pesquisa Científica, 2017). Animals were used for only one experiment and in a single behavioural test, to reduce interference from apparatus exposure. Experiments were approved by UEPA’s IACUC under protocol 06/18.

### 2.2. Sample size determination

Sample sizes were determined based on results regarding conditional approach in guppies(Pimentel et al., 2019). Time on the inspection zone was chosen as the primary endpoint, and calculations for sample sizes are valid only for that endpoint. Calculations were based on Rosner’s (2016) method for comparing two means(Rosner, 2016), and assumed α = 0.05 and power 80% on a two-tailed analysis. Based on these calculations, 15 animals were used in each group in both experiments. Animals were randomly drawn from the tank immediately before testing, and the order with which they were allocated to different conditions was randomized *via* generation of random numbers using the randomization tool in http://www.randomization.com/. Blinding was not possible, due to the positioning of the camera.

### 2.3. Experimental apparatus

The experimental apparatus was based on that described by Pimentel et al. (2019) for guppies (Pimentel et al., 2019) (Fig. 1A-B), consisting in a glass tank with 40.6 cm × 18 cm × 25 cm (l X w X h). The bottom of the tank was divided in 10 equally-sized quadrants; the quadrant that was farther from the side in which the stimulus was presented was numbered “section 1” (refuge zone), and an artificial plant was placed there to serve as refuge. Parallel to the quadrant that was farther from section 1, a computer screen (Samsung T20c310lb, 20″, LED screen, nominal brightness 20 cd/m³) allowed the presentation of animated images. These animations were produced on LibreOffice Impress (Pimentel et al., 2019), and consisted of a moving image of a leaffish (*Nandus nandus*), a sympatric predator for zebrafish (Engeszer et al., 2007; Parichy, 2015) that has been shown to induce antipredator behavior (Bass & Gerlai, 2008; Gerlai et al., 2009). Parallel to one of the longer walls of the tank (Fig. 1B), a mirror, with the length of 40.6 cm, was positioned. The mirror, therefore, occupied the whole length of the experimental apparatus and, as a consequence, created an image of a second zebrafish swimming in parallel to the focal animal. In Experiments 2 and 3, the mirror was either parallel to the tank (Cooperating mirror) or forming and angle of 45° with the tank (Defecting mirror).

**Figure 1.**
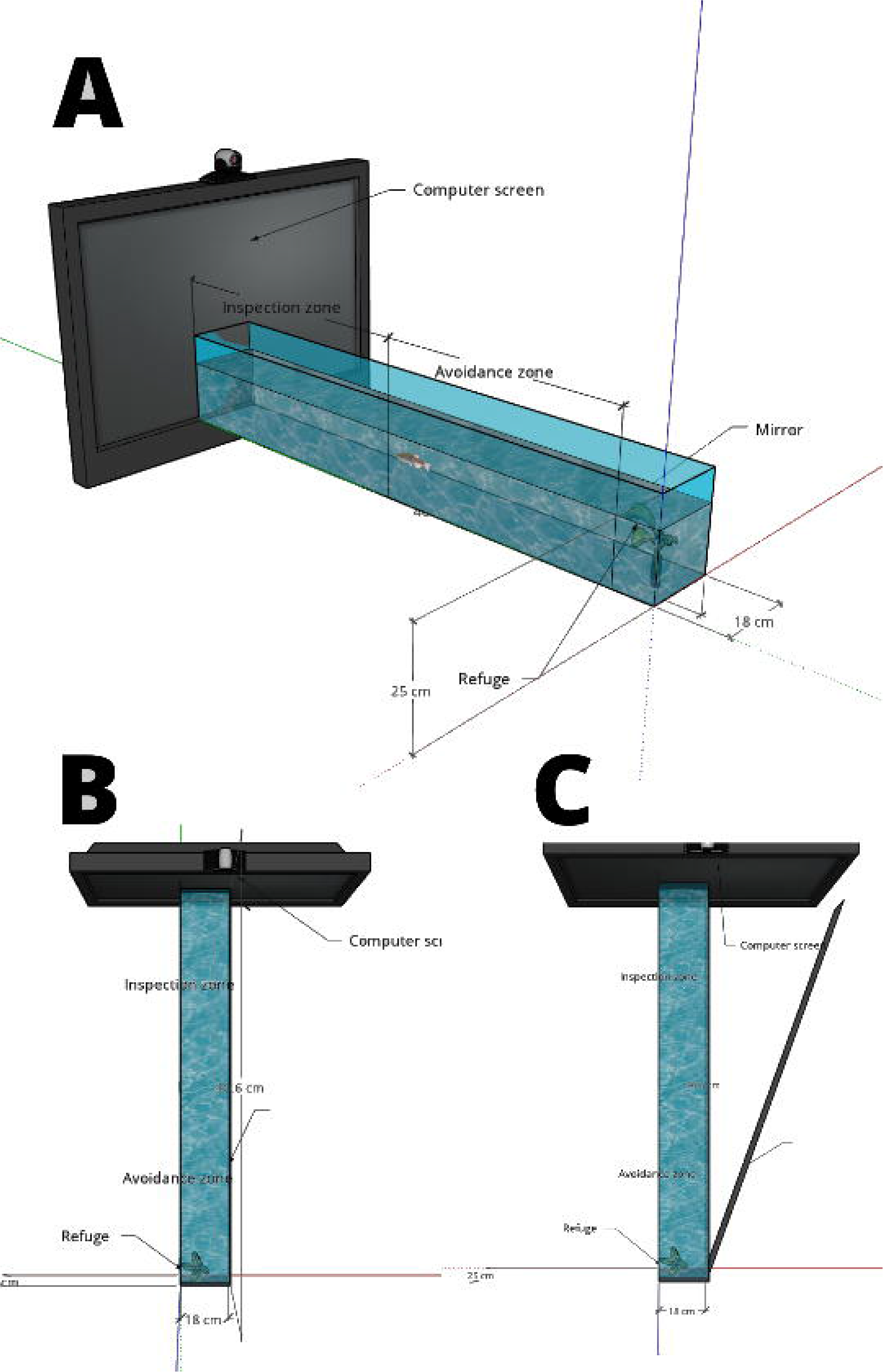
Experimental apparatus (A) Viewed from the side; (B) Viewed from above. In (B), the positions of the mirror in Experiment 2 are also indicated as CM (“Cooperating mirror”) and DM (“Defecting mirror”).

### 2.4. General design

Animals were tested individually and only once. Before beginning a trial, animals were individually transferred to the experimental tank, placed on quadrant 5 (in the center of the tank), and behaviour was filmed for 5 min while swimming freely. During this acclimation period, the screen was turned off. Animals were randomly allocated to groups, using the random number generator at https://randomization.com, to reduce allocation bias. Moreover, the order with which individuals from each group were tested was also randomized. Each trial took 15 min, and consisted in an acclimation stage (min. 0-5), immediately after the animal was introduced in the tank and during which the video was turned off, followed by the stimulation stage (min. 6-15), during which the video was turned on. Trials were recorded in digital format by a camera (Sony DCR-DVD610) mounted above the tank. Given the position of the camera in relation to the mirror, it was not possible to blind video transcription to treatment. Digital videos were later viewed, and behaviour was manually recorded using XPlo-Rat (http://scotty.ffclrp.usp.br). The following variables were recorded:

- **Time in the inspection zone:** Time spent in Sections 6 to 10 (nearest to the computer screen), expressed as percent change (time after stimulus onset – time before stimulus onset) and normalized by time block duration (i.e., time after stimulus onset was scaled to 600 s, and time before stimulus onset was scaled to 300 s);
- **Time in the avoidance zone:** Time spent in Sections 2 to 5, expressed as percent change and normalized by time block duration;
- **Time in the refuge:** Time spent in Section 1, expressed as percent change and normalized by time block duration;
- **Freezing:** Time spent immobile, except for eye and opercular movements;
- **Erratic swimming:** Number of events of zig-zagging, fast swimming episodes.

### 2.5. Experiment 1

Experiment 1 was a conceptual replication of experiments in guppies (Pimentel et al., 2019), and attempted to understand whether approach and defect behaviours were potential responses to the specific predator inspection context, or whether the animal was simply responding to the conspecific (i.e., following its own mirror image). In the “Predator” condition, the animation was shown after 5 minutes of acclimation, as described in “General design”, above; in the “No-predator” condition, no animation was shown.

### 2.6. Experiment 2

Experiment 2 was a conceptual replication of studies on cooperation vs. defection made on guppies (Dugatkin, 1988) and sticklebacks (Milinski, 1987). The rationale for the experiment is that, for two interacting fish approaching a predator, defecting (staying behind) is beneficial because the defecting individual decreases its risk but increases its payoff (e.g., information gathered) by watching the fate of the other fish. Conditional approach predicts that inspecting fish should immediately retaliate defecting conspecifics by also defecting (Bshary & Oliveira, 2015; Dugatkin, 1988; Pimentel et al., 2019). We attempt to simulate this situation by using two mirror conditions, a “cooperating” mirror (parallel to the tank) and a “defecting” mirror (in an angle of 45° to the tank); in this last situation, as the fish approached the predator, its mirror image would appear to turn away.

### 2.7. Experiment 3

Experiment 3 was an attempt to untangle shoaling tendencies from results from Experiment 2. The setup and design was exactly as in Experiment 2, except that no predator animation was shown to either mirror group.

### 2.8. Statistical analysis

For all experiments, data for time on the inspection, avoidance, and refuge zones were assessed using independent samples *t*-tests. For data on freezing and erratic swimming, one-way repeated measures analyses of variance (ANOVAs) were made, with subject as random variable. Bayes Factors (BF) were also calculated, using the ‘BayesFactor’ package in R (Morey & Rouder, 2018) (v. 0.9.12; https://richarddmorey.github.io/BayesFactor/). The prior is described by a Cauchy distribution centred around zero and with a width parameter of 0.707. This corresponds to a probability of 80% that the effect size lies between −1.25 and 1.25 for changes in time in inspection zone (prior = 0.672); between −0.49 and 0.49 for changes in time in the avoidance zone (prior = 0.385); between −0.82 and 0.82 for changes in time in the refuge zone (prior = 0.547); and between −1.36 and 1.36 for freezing (prior = 0.695). These effect sizes were based on previous work on guppies (Pimentel et al., 2019). Bayes factors were reported alongside frequentist statistics under the assumption that this can help clarify results that may be difficult to interpret using frequentist statistics alone (Malone & Coyne, 2020). *A posteriori* effect sizes were calculated as estimates of Cohen’s *d* using a bootstrap sampling distribution, with 5000 bootstrap samples, and reported with bias-corrected and accelerated confidence intervals; these calculations were made using the ‘dabestr’ package in R (Ho et al., 2018) (v. 0.2.1; https://github.com/ACCLAB/dabestr).

### 2.8. Open science practices

Experiments were not formally pre-registered. Datapackages and analysis scripts can be found on GitHub (https://github.com/lanec-unifesspa/zebrafishTFT/). Preprint versions of this paper have been submitted to bioRxiv (doi:10.1101/814434).

## 3. Results

In Experiment 1, a significant difference was found for time spent in inspection zone (Fig. 2A), with animals in the Predator condition spending more time in that area than animals in the No-Predator condition (t(17.410) = −3.065, *p* = 0.00687, *d* = 1.13 [95%CI 0.233, 1.99], BF = 10.623). No differences were found for time in the avoidance zone (Fig. 2B; t(21.219) = −1.097, *p* = 0.28746, *d* = 0.402 [95%CI −0.324, 1.17], BF = 0.546). Likewise, no differences were found for time in the refuge zone (Fig. 2C; t(26.968) = −1.291, *p* = 0.2076, *d* = 0.467 [95%CI −0.396, 1.13], BF = 0.644). Freezing was increased after animation in animals in the Predator condition compared to animals in the No-Predator condition (main effects of group: F[1, 29] = 4.606, *p* = 0.0404, BF = 1.313; main effects of time block: F[1, 29] = 4.983, *p* = 0.0445, BF = 2.239; interaction: F[1, 29] = 2.689, *p* = 0.119, BF = 3.33; *d* for Non-predator condition = 0.446 [95%CI −0.238, 1.02]; *d* for Predator condition = 0.637 [95%CI −0.207, 1.07]); while the RM-ANOVA was not significant for the interaction effect, the higher Bayes Factor suggests that the interaction model is preferred against all models.

**Figure 2.**
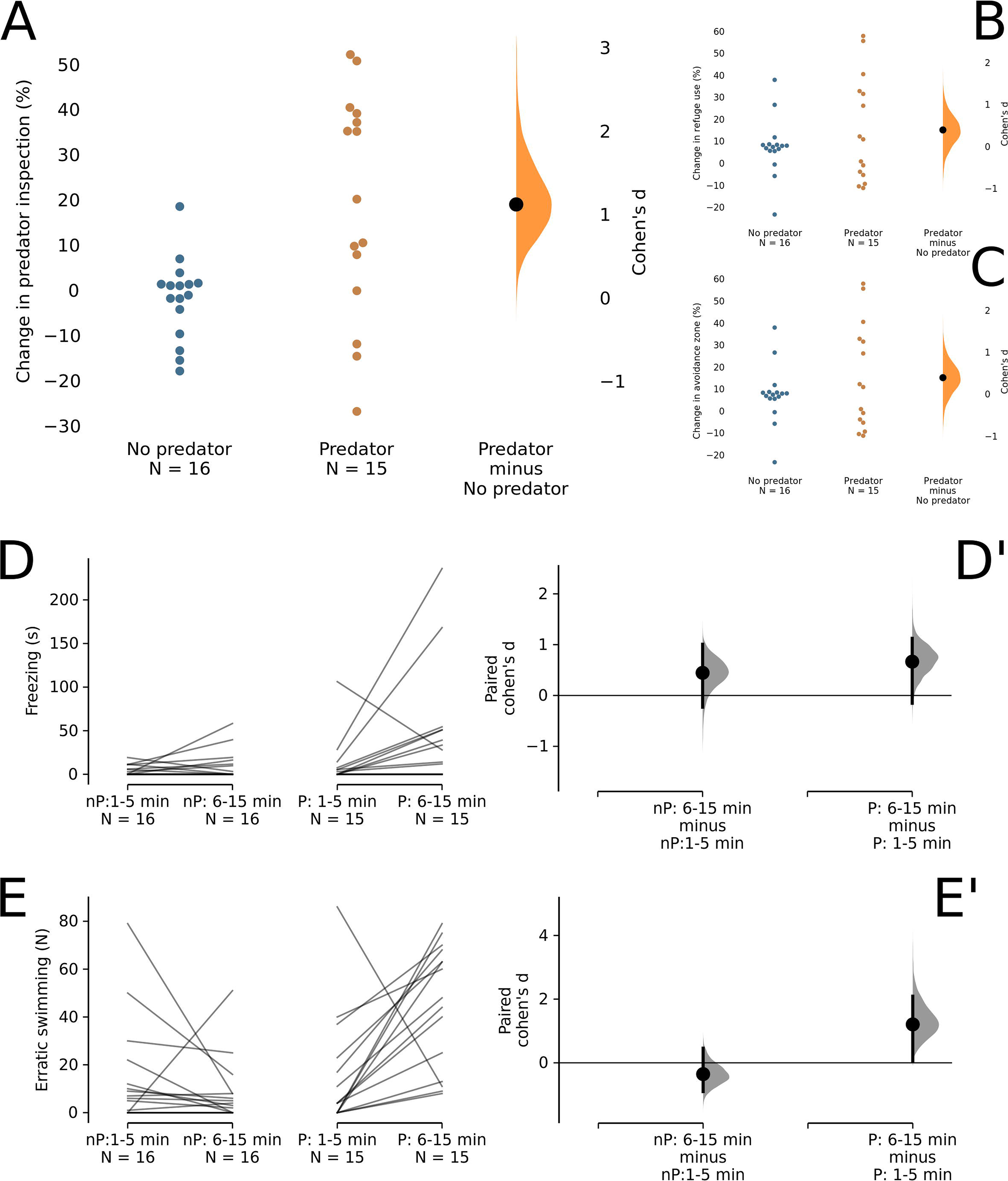
Concomitant presence of a predator animation and a cooperating mirror elicits predator approach. (A) Percent of change in the time spent in the inspection zone. (B) Percent of change in the time spent in the avoidance zone. (C) Percent of change in the time spent in the refuge zone. (D) Freezing duration. (D’) Cumming estimation plots for freezing. (E) Frequency of erratic swimming events. (E’) Cumming estimation plots for erratic swimming. In (A-C), percent changes are normalized to time block duration, individual points are shown in the left-most panel, and Cohen’s *d* between No predator and Predator groups are shown in a Gardner-Altman estimation plot. Both groups are plotted on the left axes; the mean difference is plotted on a floating axes on the right as a bootstrap sampling distribution. The mean difference is depicted as a dot; the 95% confidence interval is indicated by the ends of the vertical error bar. In (D-E), the raw data is plotted on the upper axes; each paired set of observations is connected by a line. In (D’-E’), the paired mean differences for comparisons between groups are shown in a Cumming estimation plot, with each paired mean difference plotted as a bootstrap sampling distribution of Cohen’s *d*. Mean differences are depicted as dots; the 95% confidence interval is indicated by the ends of the vertical error bar.

In Experiment 2, a significant difference was found for time spent in inspection zone (Fig. 3A), with animals in the Cooperating Mirror condition spending more time in this zone than animals in the Defecting Mirror condition (t(29) *=* −2.572, *p* = 0.0155, *d* = 0.922 [95%CI 0.187, 1.61]). Again, no differences were found for time in the avoidance zone (Fig. 3B; t(29) = −0.925, *p* = 0.3667, *d* = −0.34 [95%CI −1.17, 0.559]). Moreover, no significant differences were found for time in the refuge zone (Fig. 3C; t(29) = −0.866, *p* = 0.4, *d* = −0.311 [95%CI −0.965, 0.572]). An effect of time block (F[1, 29] = 30.126, *p* = 6.35×10-6; BF against model with subject only = 257.202) was found for freezing, but no effect of group (F[1, 29] = 0.855, *p* = 0.363; BF against model with subject only = 0.32) or interaction (F_[1, 29]_ = 1.894, p = 0.179; BF against model with subject only = 614.199) was found (paired *d* for Defecting Mirror = 0.546 [95%CI 0.139, 0.891]; paired *d* for Cooperating Mirror = 0.789 [95%CI 0.383, 1.22]). Bayes Factors suggest the superiority of a model in which time block is the main variable driving the variation in erratic swimming, An effect of time block (F[1, 30] = 10.72, *p* = 0.00268; BF against model with subject only = 39.967) was found for erratic swimming, but no effect of group (F[1, 30] = 0.113, *p* = 0.739; BF against model with subject only = 0.31) or interaction (F[1, 30] = 1.72, p = 0.2; BF against model with subject only = 9.259) was found (paired *d* for Defecting Mirror = 1.09 [95%CI 0.358, 1.67]; paired *d* for Cooperating Mirror = 1.41 [95%CI 0.851, 1.85]). Bayes Factors suggest the superiority of a model in which time block is the main variable driving the variation in erratic swimming and freezing, reinforcing the hypothesis that both cooperating and defecting mirror configurations induce fear-like behaviour.

**Figure 3.**
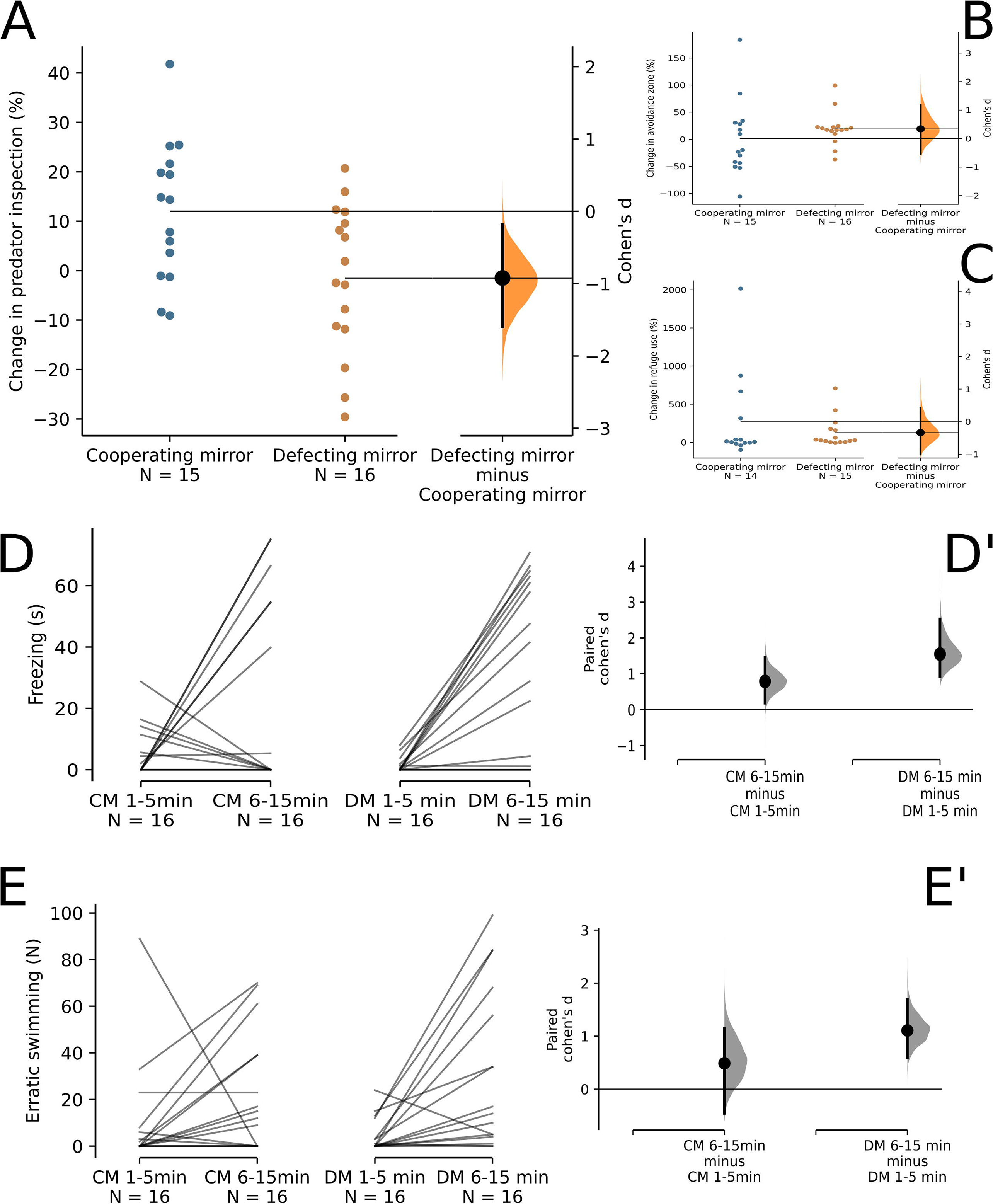
Zebrafish retaliate defections during predator inspection. (A) Percent of change in the time spent in the inspection zone. (B) Percent of change in the time spent in the avoidance zone. (C) Percent of change in the time spent in the refuge zone. (D) Freezing duration. (D’) Cumming estimation plots for freezing. (E) Frequency of erratic swimming events. (E’) Cumming estimation plots for erratic swimming. In (A-C), percent changes are normalized to time block duration, individual points are shown in the left-most panel, and Cohen’s *d* between Defecting mirror (DM) and Cooperating mirror (CM) groups are shown in a Gardner-Altman estimation plot. Both groups are plotted on the left axes; the mean difference is plotted on a floating axes on the right as a bootstrap sampling distribution. The mean difference is depicted as a dot; the 95% confidence interval is indicated by the ends of the vertical error bar. In (D-E), the raw data is plotted on the upper axes; each paired set of observations is connected by a line. In (D’-E’), the paired mean differences for comparisons between groups are shown in a Cumming estimation plot, with each paired mean difference plotted as a bootstrap sampling distribution of Cohen’s *d*. Mean differences are depicted as dots; the 95% confidence interval is indicated by the ends of the vertical error bar.

In Experiment 3, no significant differences were found between groups for time spent in inspection zone (Fig. 4A), with animals in the Cooperating Mirror condition spending a similar amount of time in this zone as animals in the Defecting Mirror condition when the animation was not presented (Fig. 4A; t_(28)_ = −0.499, p = 0.621, *d* = −0.182 [95%CI −0.898, 0.540], BF = 0.392). Again, no significant differences were found between groups for time in avoidance (Fig. 4B; t_(28)_ = 0.789, p = 0.437, *d* = 0.288 [95%CI −0.440, 1.007], BF = 0.534) and refuge (Fig. 4C; t_(28)_ = −1.064, p = 0.296, *d* = −0.389 [95%CI −1.112, 0.348], BF = 0.604). No main effects of time block (F_[1, 30]_ = 0.142, p = 0.709, BF against model with subject only = 0.271) or mirror position (F_[1, 30]_ = 0.01, p = 0.921, BF against model with subject only = 0.256) were found for freezing (Fig. 4D); no interaction effect was found as well (F_[1,31]_ = 0.191, p = 0.665, BF against model with subject only = 0.07). Finally, no main effects of time block (F_[1, 30]_ = 2.04, p = 0.164, BF against model with subject only = 0.64) nor mirror position (F_[, 30]_ = 3.019, p = 0.093, BF against model with subject only = 0.74) were found for erratic swimming (Fig. 4E); no interaction effect was found as well (F_[1, 31]_ = 1.966, p = 0.171, BF against model with subject only = 0.47). Thus, the data provide evidence against the hypothesis that, when the predator image is turned off, mirror position affects position in the tank (e.g., time in inspection or refuge zones) or fear-like responses.

**Figure 4.**
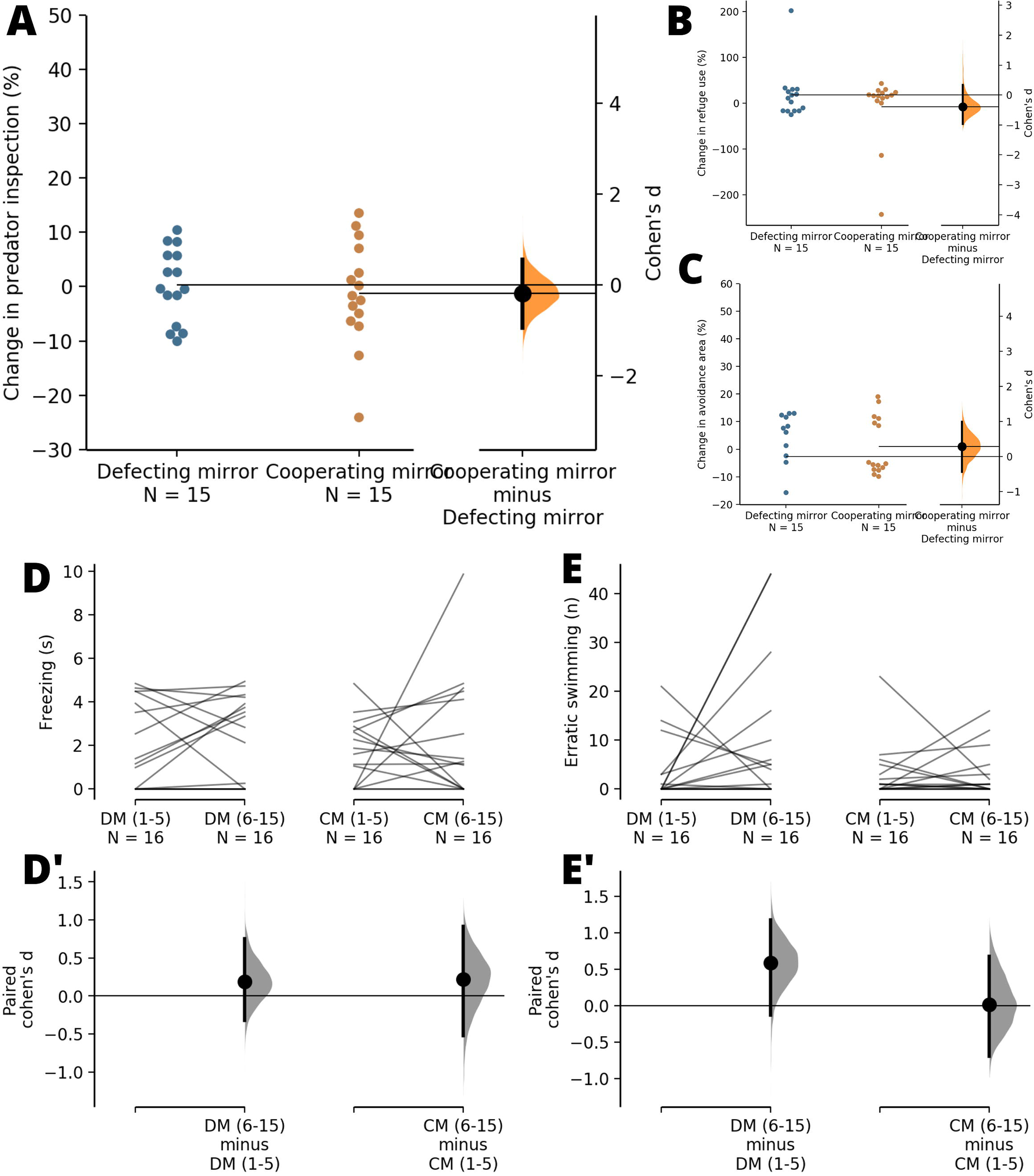
Both the mirror image and predator animation are necessary to elicit inspection behaviour and fear-like responses. (A) Percent of change in the time spent in the inspection zone. (B) Percent of change in the time spent in the avoidance zone. (C) Percent of change in the time spent in the refuge zone. (D) Freezing duration. (D’) Cumming estimation plots for freezing. (E) Frequency of erratic swimming events. (E’) Cumming estimation plots for erratic swimming. In (A-C), percent changes are normalized to time block duration, individual points are shown in the left-most panel, and Cohen’s *d* between Defecting mirror (DM) and Cooperating mirror (CM) groups are shown in a Gardner-Altman estimation plot. Both groups are plotted on the left axes; the mean difference is plotted on a floating axes on the right as a bootstrap sampling distribution. The mean difference is depicted as a dot; the 95% confidence interval is indicated by the ends of the vertical error bar. In (D-E), the raw data is plotted on the upper axes; each paired set of observations is connected by a line. In (D’-E’), the paired mean differences for comparisons between groups are shown in a Cumming estimation plot, with each paired mean difference plotted as a bootstrap sampling distribution of Cohen’s *d*. Mean differences are depicted as dots; the 95% confidence interval is indicated by the ends of the vertical error bar.

## 4. Discussion

In Experiment 1, we found that the animal spends more time in the inspection zone than in other zones when the predator stimulus is present, suggesting that the presence of a predator in the concomitant presence of a conspecific induces predator inspection, while simultaneously observing that the individual has a fear response in the form of increased erratic swimming and freezing behaviour. In experiment 2 we observed that the animal spends more time in inspection in the cooperating mirror condition than in the defecting mirror condition, and in both conditions we observed that it had fear-like behaviour. In experiment 3, we observed no changes in behaviour in any mirror position when the video was turned off, thus discarding shoaling tendencies as a driver of social behaviour under these conditions. The results of these experiments suggest that zebrafish exhibits conditional approach behaviour in situations in which also elicit fear.

In the present experiments, animals behaved towards conspecifics by approaching the mirror, and towards the predator by keeping a distance and freezing and/or displaying erratic swimming; however, when both stimuli are present, the animal emits predator inspection, entering the inspection zone and orienting towards the predator image. While it has been previously shown that visual contact with conspecifics decrease fear responses in zebrafish (Faustino et al., 2017), the presence of the conspecific image did not inhibit fear in the present experiment, as the animal still displays freezing and erratic swimming. We reported similar effects on guppies (Pimentel et al., 2019). Moreover, guppies selectively bred for high leading in a similar paradigm show no alterations in “boldness” in a simulated aerial predator task (Dimitriadou et al., 2019), suggesting that conditional approach is not related to less fear. However, field experiments show that predation risk is a factor in the evolution of predator inspection across natural guppy populations (Dugatkin & Alfieri, 1992; Edenbrow et al., 2017), suggesting that conditional approach is sensitive to threat levels.

In the cooperating mirror condition in Experiment 2, zebrafish spent more time inspecting the predator than fish in the defecting mirror condition, suggesting that zebrafish employs a “conditional approach” strategy. Similarly to “tit-for-tat”, in conditional approach the individual approaches the predator in the first movement, and subsequently approaches only if the second fish (the mirror image) swims along, retreating if the conspecific also retreated (Dugatkin, 1988, 1991, 1997). Results from our Experiment 2 suggest that the animal retaliates after defection (i.e., reduces inspection in the defecting mirror condition).

Zebrafish were thus shown to be able to change their strategy from avoiding to approaching predators when conspecifics are present. If in visual contact with a conspecific (their own reflection), they proceed to spend more time inspecting predator (cooperation mirror condition), a situation that significantly changed in the defecting mirror condition, as their company was absent. This looks like a typical “conditional approach” situation: conditionally, the individual swims towards the predator in the first movement, but subsequent close inspection only occurs if the second fish (the mirror image) swims along, retreating immediately if the accompanying conspecific also abandons (Dugatkin, 1988). This is an indication of the social conditions necessary the evolution of predator approach behaviour as a cooperative strategy in zebrafish, as in guppies and sticklebacks (Brosnan et al., 2003; Croft et al., 2006; Dugatkin & Alfieri, 1991a; Edenbrow et al., 2017; Huntingford et al., 1994): strong social bonds between conspecifics that enable their first risky move (cooperative behaviour) which can confer benefits to those inspecting and to the shoal (Bshary & Oliveira, 2015; Croft et al., 2006; Dugatkin, 2013). However, if facing defecting shoal-mates, individuals may retaliate, ceasing inspection but are able to “forgive” if baseline conditions are restored. Our findings allow us to include zebrafish in the restrict group of teleost fish species capable of complex social decision making, responding to the quality of social bonds by being able to change strategy as conditions change.

Currently, the benefits for the inspecting zebrafish are unknown. Reciprocal altruism suggests that the benefit of any cooperative action involve increased probability of reciprocation in the future (Trivers, 1971) – that is, an individual would be incentivized to inspect a predator if a shoal-mate is more likely to inspect in the future (Edenbrow et al., 2017). Reciprocal altruism can also evolve by generalized reciprocity, in which cooperation evolves in small groups when individuals base their decision on the outcome of previous interactions with anonymous partners (i.e., without the need for individual recognition and specific social memory) (Pfeiffer et al., 2005). Generalized reciprocity is thought to be favoured by contexts in which cooperation is more advantageous in a cooperative than in a noncooperative environment, leading to stable levels of cooperation when the decision to stay or leave a group co-evolves with the decision to cooperate (I. M. Hamilton & Taborsky, 2005). Indeed, both Experiments 1 and 2 suggest that, as zebrafish are more likely to inspect when a “shoal-mate” (mirror image) is present, and when the shoal-mate reciprocates inspection (i.e., in the “cooperating”, but not in the “defecting” mirror). This does not discard, of course, other benefits of inspecting, although currently these have not been investigated in zebrafish. Indeed, our results are also consistent with generalized reciprocity, in which simple decision rules such as “defect when encountering noncooperation” is sufficient to lead to stable cooperation levels (Aktipis, 2004). In guppies, cooperative animals (i.e., animals with high levels of predator inspection) are preferred by conspecifics (Dugatkin & Alfieri, 1991a; Godin & Dugatkin, 1996), and assortative interactions and social selection are expected to play a role (Edenbrow et al., 2017). It has been previously suggested that conditional approach is better understood in a “social competence” approach (Bshary & Oliveira, 2015; Taborsky & Oliveira, 2012), and therefore does not require some form of kin or group selection to be at work (Pimentel et al., 2019). In this context, situational cues (such as presence or absence of females, or varying risk levels), together with other factors, allow individuals to assess the situation and make behavioural decisions.

Zebrafish has been previously shown to display predator inspection, at low levels, without the presence of conspecifics (Dugatkin et al., 2010; Pannia et al., 2014). Our results are the first demonstration that predator inspection is related to cooperative-like behaviour in zebrafish, even when the animal displays fear-like behaviour towards the predator. These results suggest that the social behaviour of zebrafish is more complex than previously thought, and can be further exploited in the study of the neural bases of altruism in an interesting model organism.

## Acknowledgements

AFP was the recipient of a CNPq studentship.

